# Mass spectrometry and NMR spectroscopy profiles of red and pink Rhododendron flower petals establish them as rich sources of bioactive secondary metabolites

**DOI:** 10.1101/2023.12.26.573342

**Authors:** Shagun Shagun, Maneesh Lingwan, Shyam Kumar Masakapalli

**Affiliations:** School of Biosciences and Bioengineering, Indian Institute of Technology Mandi, Kamand 175075, India

**Author notes:** Address correspondence to Shyam Kumar Masakapalli.

**Keywords:** *Rhododendron arboreum*, *Rhododendron campanulatum*, comprehensive phytochemical profiling, multi-analytical platforms, bioactive metabolites

## Abstract

Rhododendron petals are considered high-value owing to their commercial utility, national/state flower status in certain countries, and bioactive potential from recent studies. Profiling and quantitative analysis of the bioactive metabolites would evaluate if they can be natural sources. This study is focused on comprehensive profiling of secondary metabolites in the petals of Red and Pink Rhododendron flowers (*R. arboreum* and *R. campanulatum*) using Mass Spectrometry (GC-MS, LC-MS/MS) and Proton-Nuclear Magnetic Resonance (^1^H-NMR) Spectroscopy. The profiling highlighted the presence of secondary metabolites belonging to phenolic acids and flavonoids. Specifically, the flowers are rich in promising bioactive molecules such as quinic acid, chlorogenic acid (3-O-caffeoyl quinic acid), protocatechuic acid, coumaroyl quinic acids, catechin, epigallocatechin, and shikimic acid. The profiles are correlated with the metabolic pathways which reflected the activity of shikimic acid, phenolic acid and flavonoid biosynthetic pathways. These metabolites are well reported for their bioactive potential as anti-oxidative, anti-viral, anti-cancerous, anti-diabetic, anti-inflammatory, etc. While the quantitative and multivariate analysis showed variations in the levels of phenolic acids and flavonoids, it is established that red and pink Rhododendron flower petals are a rich source of bioactive phytochemicals of interest to the phytochemical Industry.

## 1. Introduction

Indian Himalayan Region (IHR) is renowned for its rich diversity of medicinal plant species due to its varied climatic conditions. Most of these species are endemic to some regions in the Himalayas (Shagun & Masakapalli, 2023). Rhododendron is a genus of flowering plants mainly distributed in higher elevations and known to have diverse medicinal and commercial uses (Kumar et al., 2019). Around 1,000 species of this genera are distributed worldwide, out of which 87 species are present in the IHR. The *Rhododendron arboreum* and *Rhododendron campanulatum* are among the six species that are found to be present abundantly in the western Himalaya region (Madhvi et al., 2019; Sekar et al., 2010). *R. arboreum* (red Rhododendron) has been recognised as national flower of Nepal, state flower of Nagaland and state tree of Uttarakhand, India whereas *R. campanulatum* (pink Rhododendron) is state flower of Himachal Pradesh (Keshari et al., 2017; Gautam et al., 2020; Thakur et al., 2023). For the local people, Rhododendrons are a sustainable source of income by providing various services such as beverages, food supplements, fuelwood, and other medicinal benefits. Different parts of these plants exhibit many ethnobotanical uses and have been used for the treatment of many health-related problems (Singh et al., 2020).

*R. arboreum* has red-colored petals and are consumed by local inhabitants in various forms, such as juices, jams, pickles, and chutney (Gautam et al., 2020; Kumar et al., 2019). Traditionally, these red petals in various forms (fresh, dried, and juice) are used for the treatment of various ailments like diarrhea, blood dysentery, high altitude sickness, headaches, diabetes, rheumatism, etc (Bhatt et al., 2022; Lingwan et al., 2021). The young leaves of *R. arboreum* have medicinal properties and are applied on the forehead to relieve headaches but cause intoxication in large quantities (Watt, 1982). The phytochemical profiling of *R. arboreum* flower petals indicated the potential presence of various phytochemicals such as quinic acid, chlorogenic acid, coumaroyl quinic acid, catechin, p-coumaric acid, etc., in the hot aqueous extract which has been reported to have potential antiviral and other health beneficial properties (Lingwan et al., 2021). Recently, Bhatt et al., 2022 identified anthocyanins (cyanidin-3-O-β-galactoside, cyanidin-3-O-α-arabinoside and cyanidin-3-O-rhamnoside), phenolic acids and flavonoids in *R. arboreum* flower petals.

*R. campanulatum* species (pink-colored flowers) have many ethnobotanical uses and have been used for long. Due to its overexploitation for medicinal uses and other factors like baseline shifting and flowering pattern changes, this species has been enlisted as endangered in IUCN Red data list (Thakur et al., 2023). The bark of this species is used to treat jaundice, piles, and liver disorders (Singh et al., 2020). The roots of this species are used for the alleviation of various diseases such as cough, headache, boils, and fever (Bartwal et al., 2020). The leaves and flowers of *R. campanulatum* are considered to be toxic but traditionally they are used as a remedy for various ailments. The leaves, when mixed with tobacco, are helpful in the treatment of chronic rheumatism, syphilis, cold, hemicranias, and sciatica (Bartwal et al., 2020; Prakash et al., 2016), and the flowers are used for skin diseases, throat pain, and body ache (Kunwar et al., 2010). The ethnobotanical uses of this species are known so far, but the phytochemical profiling of this species is yet to be studied elaborately, which is the main focus of this study. The comprehensive phytochemical profiling of the *R. arboreum* and *R. campanulatum* petals hot aqueous extract was done by using multi-analytical platforms. For qualitative and quantitative analysis, the extracts were subjected to LC-MS/MS (Liquid Chromatography-Mass Spectrometry), GC-MS (Gas Chromatography-Mass Spectrometry), and ^1^H-NMR spectroscopy. The qualitative analysis highlighted the presence of various bioactive secondary metabolites including phenolic acids (quinic acid, chlorogenic acid, protocatechuic acid, p-coumaric acid, coumaroyl quinic acids), and flavonoids (catechin, epigallocatechin). The quantitative and multivariate analysis showed variations in the abundances of secondary metabolites among both the flower species. This work is a ready reference for the phytochemical profiles of the Rhododendron petals.

## 2. Material and methods

### 2.1. Plant material collection and processing

*Rhododendron arboreum* and *Rhododendron campanulatum* flowers were collected from the hills in the mid-Himalayan region [>1500 m height above mean sea level (AMSL)] in district Mandi, Himachal Pradesh, India. The red *R. arboreum* flowers were collected from the Parashar Lake area, while the pink *R. campanulatum* blossoms were collected from the Diana Park area. The petals were removed from the flowers and thoroughly cleaned with water. The washed petals were vacuum-dried before being pulverized to fine powder. For subsequent analysis, the powdered petals were kept in airtight containers at room temperature.

### 2.2. Phytochemicals extraction

*R. arboreum* and *R. campanulatum* phytochemicals extracts were prepared by boiling the powdered petals (50 mg DW in 1 ml distilled water) at 70 °C for 5 min with continuous shaking at 500 rpm. The extract was centrifuged at 13,000 rpm for 10 minutes at room temperature to pellet down any insoluble material. The supernatant was then filtered through syringe filters with pore size 0.22 μm and transferred to a new microcentrifuge tube further for Mass Spectrometry (LC-MS/MS & GC-MS) and Proton Nuclear Magnetic Resonance (^1^H NMR) Spectroscopy analysis (Lingwan et al., 2021).

### 2.3. Phytochemical characterization of the extracts

#### 2.3.1. Liquid Chromatography-Mass Spectrometry (LC-MS/MS) based characterization

The suitable filtered extracts were transferred to the glass vials for LC-MS/MS characterization. Different concentrations (0.2-1 mg/ml) of quinic acid and chlorogenic acid standards were prepared in distilled water. The prepared standards were filtered through syringe filters with pore size 0.22 μm and transferred to glass vials. The Rhododendron petals phytochemicals and standards were characterized by LC-ESI-QTOF-MS/MS instrument with a UHPLC equipped with QTOF MS/MS. The C-18 column was used for the separation of the various phytochemicals. Column temperature was set at 30 °C. The mobile phase used was (A) water with formic acid (0.1%) and acetonitrile (B). The gradient program used as per solvent B is a linear gradient of B from 7-15% for 0-3min, 15-20% for 3-5min, isocratic elution of 20% B for 5-8min and further linear gradient from 20-25% B for 8-10 min. The flow rate used was 0.2 ml/min. The injection volume was 5 μl (Tian et al., 2020).

#### 2.3.2. Gas Chromatography-Mass Spectrometry (GC-MS) based characterization

For GC-MS analysis, 50 μl filtered extract was taken, and ribitol (0.01% w/v) was added as an internal standard. This mixture was dried in the speed vacuum. The dried extract of *R. arboreum* and *R. campanulatum* petals were derivatized by MSTFA. To obtain the phytochemicals derivatives, 50 μl pyridine with MeOX (20 mg/ml) was added and incubated at 37 °C for 2 hours with shaking at 900 rpm. Then 70 μl MSTFA was added to the samples and incubated at 37 °C for 30 minutes with shaking at 900 rpm. The derivatized samples were centrifuged at 13,000 rpm for 10 minutes at room temperature, and the supernatant was transferred to GC-MS vials. GC-MS data was acquired in splitless mode on a GC (Agilent instrument) equipped with an HP-5MS column (Agilent 19091S-433; 30 m x 250 μm x 0.25 μm). The carrier gas (helium) flow rate was maintained at 0.6 ml/min, and the injection volume was 1 μl. The initial oven temperature was set from 50 °C to 70 °C with 5 minutes hold time. Further increased to 200 °C at 10 °C/min with 10 minutes hold time and final ramped to 300 °C temperature at 5 °C/min and a hold time of 10 min. The total run time was 60 min. The raw GC-MS spectra were baseline-corrected by using Metalign software (Lommen et al., 2012). The phytochemicals were identified using library NIST 17 (National Institute of Standards and Technology, Maryland) and further confirmed with commercially available standards. Retention time, m/z, and peak area were retrieved for each phytochemical (Masakapalli et al., 2014).

#### 2.3.3. ^1^H NMR spectroscopy-based analysis of the extracts

The filtered aliquots (500 μl) of the extracts were dried in a speed vacuum. The dried extracts were redissolved in deuterated water (D2O) containing 0.01% w/v 4,4-dimethyl-4-silapentane-1-sulfonic acid (DSS, internal standard). The samples were mixed adequately by vortexing for 5 minutes and then transferred to the NMR tubes (5mm diameter). Different concentrations (0.1-2 mg/ml) of Quinic acid and Chlorogenic acid standards (prepared in deuterated water) were also subjected to ^1^H-NMR spectroscopy to confirm and quantify in the extract samples. The ^1^H-NMR spectroscopy of the extract was performed on a JEOL JNM ECX-500 NMR instrument. 64 scans with a 5-second relaxation delay were used to record the spectra of the extract samples. The spectra were manually processed for phasing, baseline correction, and referencing (DSS at 0 ppm). Further, the normalization of each spectrum was done with reference to the internal standard, and the relative area was taken further for the quantification of quinic acid and chlorogenic acid.

### 2.4. Statistical data analysis, Multivariate data analysis, and Pathway mapping

Results were presented as mean ± standard deviation (SD). Graph pad 8.0 was used for the statistical data analysis [t-test and analysis of variance (ANOVA)] at a significance level of *p* < 0.05. MetaboAnalyst 5.0, an online tool, was used for the multivariate statistical analysis based on the respective abundances of all the phytochemicals and secondary metabolites in the samples. Data pre-processing steps were performed, including data normalization, transformation, and scaling. Data normalization was done by using normalization by sum. Log transformation and Pareto scaling were used for data transformation and data scaling. Further Principal component analysis (PCA), a Heat map, and Variable Importance in Projection (VIP) plots were generated and analyzed to see the variations among both species (Pang et al., 2021). Further, the Pathway was mapped for all the phenolic acids and flavonoids present in the Rhododendron species (*R. arboreum* and *R. campanulatum*) petals.

## 3. Results and Discussion

### 3.1. Qualitative analysis using multi-analytical platforms-based profiling highlighted the presence of promising bioactive metabolites in Rhododendron flower petal extract

*R. arboreum* and *R. campanulatum* petals hot aqueous extracts were subjected to mass spectrometry (LC-MS/MS and GC-MS) and ^1^H-NMR to identify the various phytochemicals. The profiling highlighted the presence of different secondary metabolites, organic acids, and a few unknown metabolites **(Figures 1 & 2, Table 1, Supplementary Figure 1, and Supplementary Tables 1-3)**. Specifically, the flowers are rich in promising bioactive molecules such as quinic acid, chlorogenic acid (3-O-caffeoyl quinic acid), protocatechuic acid, coumaroyl quinic acids, Catechin, epigallocatechin and shikimic acid **(Figure 1 & 2, Table 1 and Supplementary tables 1-3)**. Further, the quinic acid and chlorogenic acid were confirmed by comparing with analysis of commercially available standards **(Supplementary figures 2-4)**.

**Table 1:** LC-ESI-QTOF-MS/MS characteristics of the secondary metabolites identified from *Rhododendron arboreum* and *Rhododendron campanulatum* petals hot aqueous extract.

| S.No. | Metabolites Identified | Retention time | Chemical formula | [M-H]<sup>-</sup> m/z | Fragment ions m/z (MS/MS) | Metabolites class |
| --- | --- | --- | --- | --- | --- | --- |
| 1 | Quinic acid | 1.8 | C<sub>7</sub>H<sub>12</sub>O<sub>6</sub> | 191 | MS<sup>2</sup> [191]: 108, 109, 127 | Phenolic acid |
| 2 | Chlorogenic acid | 2.8 | C<sub>16</sub>H<sub>18</sub>O<sub>9</sub> | 353 | MS<sup>2</sup> [353]: 191 | Phenolic acid |
| 3 | Protocatechuic acid | 3.0 | C<sub>7</sub>H<sub>6</sub>O<sub>4</sub> | 153 | MS<sup>2</sup> [153]: 108, 109 | Phenolic acid |
| 4 | Catechin | 7.4 | C<sub>15</sub>H<sub>14</sub>O<sub>6</sub> | 289 | MS<sup>2</sup> [289]: 123, 109, 151 | Flavonoid |
| 5 | Coumaroyl quinic acid-1 | 8.6 | C<sub>16</sub>H<sub>18</sub>O<sub>8</sub> | 337 | MS<sup>2</sup> [337]: 191, 119 | Phenolic acid |
| 6 | Coumaroyl quinic acid-2 | 9.7 | C<sub>16</sub>H<sub>18</sub>O<sub>8</sub> | 337 | MS<sup>2</sup> [337]: 191, 119 | Phenolic acid |

**Figure 1:**
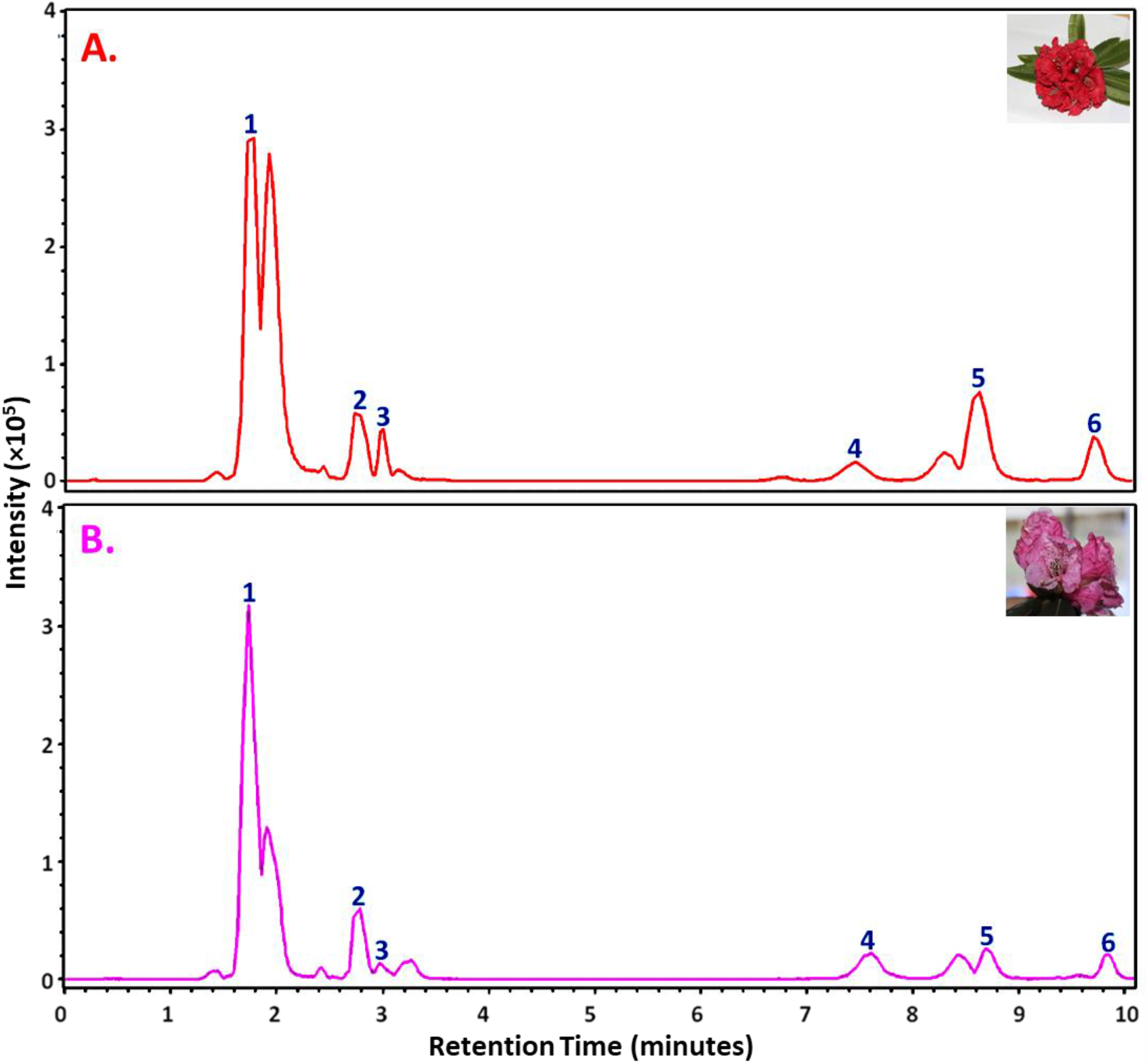
LC-MS/MS chromatograms of **A**. Rhododendron arboreum **B**. *Rhododendron campanulatum* petals hot aqueous extract detected in negative ion mode. Secondary metabolites identified are quinic acid (1), chlorogenic acid (2), protocatechuic acid (3), catechin (4), and coumaroyl quinic acid (5 & 6). Quinic acid and chlorogenic acid retention times were confirmed with the analysis of commercially available known standards (Supplementary figure 1&2).

**Figure 2:**
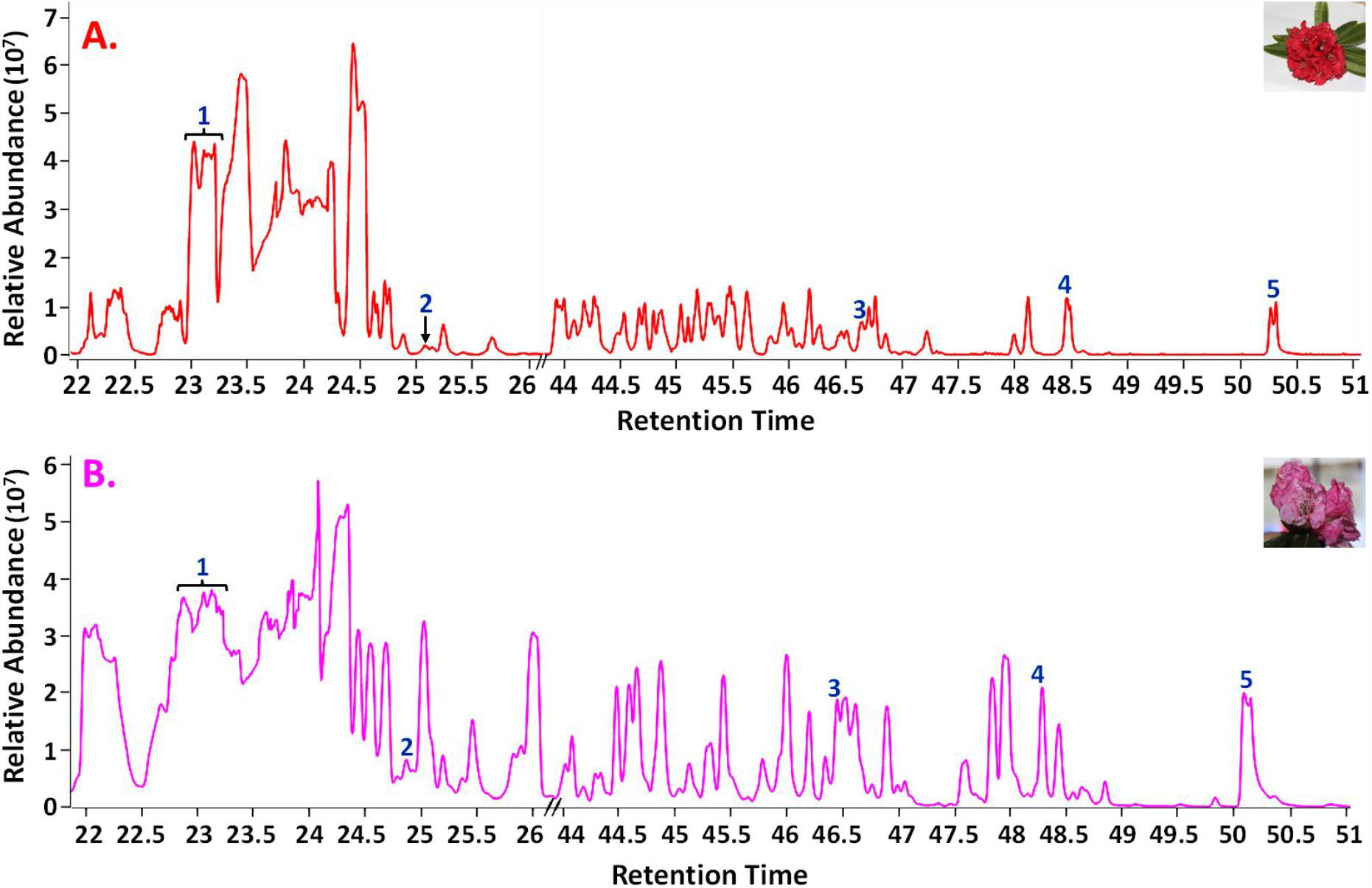
GC-MS chromatograms of hot aqueous extracts of **A**. *Rhododendron arboreum* and **B**. *Rhododendron campanulatum* petals. The secondary metabolites identified are 1. Quinic acid, 2. p-coumaric acid, 3 & 4. 5-O-Coumaroyl-D-quinic acid, and 5. Chlorogenic acid.

#### 3.1.1. LC-ESI-QTOF-MS/MS-based characterization confirmed the presence of phenolic acids and flavonoids in Rhododendron petals extract

The phytochemicals from *R. arboreum* and *R. campanulatum* petals were characterized using LC-ESI-QTOF-MS/MS and compared. Negative ionization mode with MS scan mode from 80 to 1300 m/z was applied for the phytochemical characterization in hot aqueous extract of petals. Further, the metabolites were identified by comparing the phytochemicals m/z and fragmentation patterns with known standards available and from the literature. A total number of 11 phytochemicals were detected from the petal’s hot aqueous extract. The identified metabolites belong to different metabolite classes, such as phenolic acids, flavonoids, organic acids, and a few unknowns **(Figure 1, Table 1, Supplementary Figure 1, and Supplementary Table 1)**. The major phenolic acids and flavonoids identified are quinic acid, chlorogenic acid, protocatechuic acid, catechin, and coumaroyl quinic acids **(Figure 1 and Table 1)**. The presence of quinic acid (retention time -1.8 minutes) and chlorogenic acid (retention time -2.8 minutes) was confirmed after comparing with known standards available **(Supplementary figure 2&3)**.

**Table 1** shows the various phenolic acids and flavonoids identified from the *R. arboreum* and *R. campanulatum* petals hot water extract based on their characteristic precursor ion [M-H]**-** m/z and MS/MS fragments. Five phenolic acids and one flavonoid [metabolites 1-6] were identified **(Table 1)**. Metabolite 1 with characteristic precursor ion [M-H]-m/z at 191 was identified as Quinic acid with MS/MS fragments ions m/z at 108, 127, 109. Metabolite 2 was characterized as Chlorogenic acid with an observed precursor ion [M-H]-m/z at 353 and MS/MS fragment ion m/z at 191. The presence of Quinic and Chlorogenic acid was confirmed by comparing the precursor ion [M-H]-m/z and MS/MS fragments ions with the available standards. Based on the characteristic precursor ion [M-H]-m/z at 153 and MS/MS fragments ions at 108 and 109, the phytochemical 3 was identified as Protocatechuic acid with the characteristic precursor ion [M-H]-m/z at 153. The MS/MS fragment ions m/z at 108 and 109 are consistent with the m/z reported in the literature for the same metabolite (Bouhafsoun et al., 2018). Phytochemicals 5 and 6 were with precursor ions [M-H]-m/z at 337 exhibiting fragment ions at m/z at 191 and 119 were categorized as Coumaroyl quinic acid. The presence of coumaroyl quinic acids was confirmed by comparing the precursor ions [M-H]- and fragment ions at m/z with the reported literature (Yang et al., 2022). One flavonoid, i.e., Catechin (phytochemical 4), was identified with precursor ion [M-H]-m/z at 289. The MS/MS fragment ions m/z at 123, 109, and 151 are consistent with the already reported m/z for Catechin (Singh et al., 2018).

Previously, Bhatt et al., 2022, reported the presence of different phenolic acids, flavonoids, and anthocyanins in the acidified methanolic extract of *R. arboreum* petals by using HPLC. Various proanthocyanidins, glycosides, and hydroxycinnamates were detected from the methanolic extracts of leaves from 14 Rhododendron species using LC-MS (Jaiswal et al., 2012).

#### 3.1.2. GC-MS analysis showed the presence of phenolic acids in Rhododendron hot aqueous petals extract

The hot aqueous extracts of *R. arboreum* and *R. campanulatum* petals were analyzed through GC-MS, which showed the presence of various metabolites. These phytochemicals belong to various classes, such as phenolic acids, flavonoids, organic acids, and a few intermediates **(Figure 2, Supplementary Tables 2 and 3)**. Twenty-seven phytochemicals were identified from the hot aqueous extract of *R. arboreum* petals. Among these, a few secondary metabolites were identified, which include protocatechuic acid, quinic acid, p-coumaric acid, 5-O-coumaroyl-D-quinic acid, chlorogenic acid, shikimic acid, and catechin **(Figure 2A, Supplementary table 2)**. From *R. campanulatum* petals hot aqueous extract, a total of thirty-seven phytochemicals were identified. Major secondary metabolites identified were quinic acid, p-coumaric acid, 5-O-coumaroyl-D-quinic acid, 3-O-coumaroyl-D-quinic acid, chlorogenic acid, shikimic acid, catechin and epigallocatechin **(Figure 2B, Supplementary table 3)**. Several phytochemicals identified in the red petals of Rhododendron matches with that of earlier studies except for previously misannotated 5-O-Feruloyl-quinic acid (Lingwan et al 2021). Analysis with standards and further validation with LC-MS/MS established the presence of 3-O-Caffeoyl-quinic acid (chlorogenic acid) in this study.

#### 3.1.3. ^1^H-NMR analysis confirmed the presence of quinic acid and chlorogenic acid in Rhododendron petal’s hot aqueous extract

The hot aqueous petal extract of *R. arboreum* and *R. campanulatum* was analyzed using ^1^H-NMR. Also, the quinic acid and chlorogenic acid standards were run (at different concentrations) for their confirmation in the flower petals’ hot aqueous extract **(Supplementary Figures 4 & 5)**. The ^1^H-NMR analysis confirmed the presence of quinic acid **(Supplementary Figure 4)** and chlorogenic acid **(Supplementary Figure 5)** in the extract of both species’ hot aqueous petals. Previously, Lingwan et al., 2021 confirmed the presence of quinic acid in the *R. arboreum* hot aqueous petals extract by using ^1^H-NMR. Characterization of two significant anthocyanins, i.e., cyanidin-3-O-β-galactoside and cyanidin-3-O-α-arabinoside, in the *R. arboreum* extract was done by using 1D and 2D NMR analysis in a study carried out by Bhatt et al., 2022.

### 3.2. Multi-analytical platforms-based quantitative analysis showed significant variations in the abundance of secondary metabolites among Rhododendron species

The relative/absolute abundances (expressed as mean ± standard deviation) of the various metabolites profiled from *R. arboreum* and *R. campanulatum* petals hot aqueous extract by using multi-analytical platforms were compared. The statistical analysis showed significant variations among the levels of various profiled phenolic acids, flavonoids, and organic acids **(Figures 3-5, Supplementary Figure 6)**.

**Figure 3:**
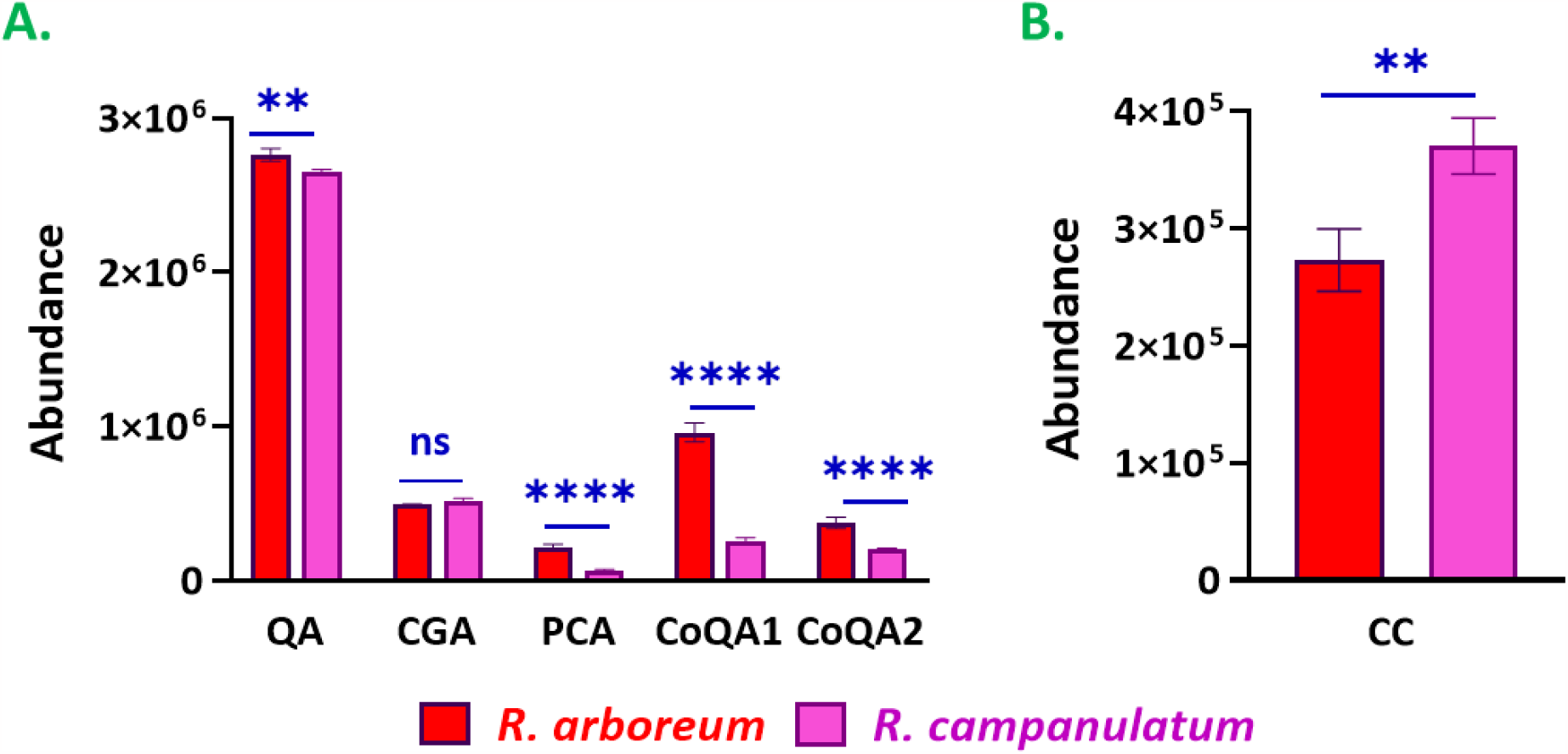
The difference in the abundances of **A**. Phenolic acids and **B**. Flavonoid [expressed as Mean ± Standard deviation (n=3)] characterized through LC-MS/MS in *Rhododendron arboreum* and *Rhododendron campanulatum* petals hot aqueous extracts showed the significant differences (at a significance level of *p* < 0.05) in their levels among both the species. (Abbreviations used: QA-Quinic acid; CGA-Chlorogenic acid; PCA-Protocatechuic acid; CoQA-Coumaroyl quinic acid; CC-Catechin, ** means p < 0.05 **** means p < 0.0001, ns-not significant)

#### 3.2.1. LC-ESI-QTOF-MS/MS-based profiling showed significant variations in the abundance of secondary metabolites among Rhododendron species

The abundances [expressed as Mean ± Standard deviation (n=3)] of the various secondary metabolites from *R. arboreum* and *R. campanulatum* petals hot aqueous extract by using LC-ESI-QTOF-MS/MS were compared. The statistical analysis showed significant differences (at a significance level of *p* < 0.05) in their levels in both species **(Figure 3)**. Among the identified phenolic acids, abundances of quinic acid (1-fold), protocatechuic acid (3.4-fold), caffeoylquinic acids-1 (3.7-fold), and caffeoylquinic acids-2 (1.9-fold) were found to be significantly higher in *R. arboreum* petals extract. The abundance of chlorogenic acid was 1.1-fold higher in *R. campanulatum* petals extract than *in R. arboreum; h*owever, the difference was not statistically significant **(Figure 3A)**. Significant differences were observed among the flavonoid catechin (CC) levels in both species. Catechin was 1.4-fold higher in *R. campanulatum* than *in R. arboreum* (**Figure 3B)**.

#### 3.2.2. ^1^H-NMR analysis showed significant variations in the absolute abundances of quinic acid and chlorogenic acid among Rhododendron species

The quinic acid and chlorogenic acid were quantified in the hot aqueous petals extract of *R. arboreum* and *R. campanulatum* using ^1^H-NMR. The quantitative analysis of quinic acid and chlorogenic acid showed statistically significant differences (at a significance level of *p* < 0.05) between both the species petals hot aqueous extract **(Figure 4)**. Quinic acid was found to be present at a 1.3-fold higher concentration in hot aqueous petals extract in *R. arboreum* than in *R. campanulatum* **(Figure 4A)**. At the same time, the abundance of Chlorogenic acid was found to be 1.5-fold higher in *R. campanulatum* than *in R. arboreum* **(Figure 4B)**.

**Figure 4:**
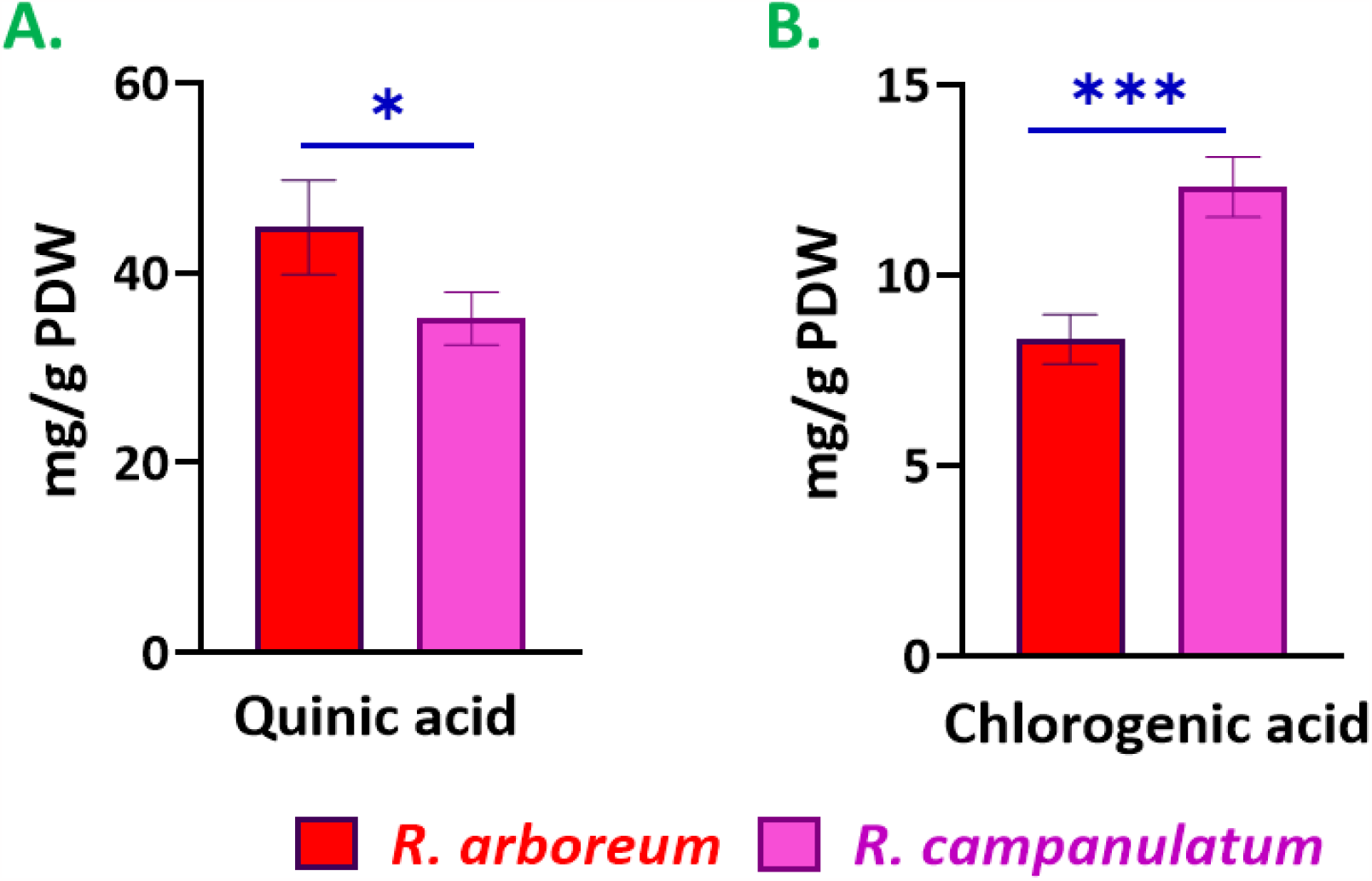
Relative absolute abundances expressed in mg/g DW (Dry Weight) of **A**. Quinic acid and **B**. Chlorogenic acid quantified from *Rhododendron arboreum* and *Rhododendron campanulatum* petals hot aqueous extracts. The data was presented as mean ± standard deviation (n=4), which showed significant differences (at a significance level of p < 0.05) in their levels among both species.

### 3.3. Multivariate statistical analysis showed the variations among Rhododendron (*R. arboreum* and *R. campanulatum*) species

Multivariate statistical analysis captured the variations in the secondary metabolites profiling of Rhododendron species **(Supplementary Figure 7)**. PCA (Principal Component Analysis) plot indicated the distinct clusters of *R. arboreum* and *R. campanulatum* species, where principal components (PCs) account for variance, i.e., 91.6% (PC1) and 3.3% (PC2) in the phytochemical’s profiles of both the species **(Figure 7A)**. Further, the VIP plot was generated, which showed that protocatechuic acid, quinic acid, and shikimic acid are the key phenolic acids responsible for maximum variations in both species (**Figure 7B)**.

### 3.4. *Rhododendron* (*R. arboreum and R. campanulatum*) petals are rich in bioactive metabolites

The biosynthetic pathway was plotted for the secondary metabolites detected in Rhododendron (*R. arboreum & R. campanulatum*) species through the multi-analytical platform **(Figure 5)**. The pathways analysis showed the completeness of the shikimic acid pathway, phenolic acids, and flavonoids biosynthetic pathways, which lead to the biosynthesis of shikimic acid, different phenolic acids (quinic acid, protocatechuic acid, p-coumaric acid, chlorogenic acid, and coumaroyl quinic acids) and flavonoids (catechin and epigallocatechin). The identified phenolic acids and flavonoids are reported to have various health-beneficial properties, such as anti-viral (Lyu et al., 2005; Quiroz et al., 2014; Zanello et al., 2015; Dai et al., 2017; Ding et al., 2017; Kwon et al., 2020), anti-cancerous (Xue et al., 2017; Singh et al., 2018; Changizi et al., 2021), anti-diabetic (Samarghandian et al., 2017; Singh et al., 2021), anti-inflammatory and anti-oxidative (Jian-Guo et al., 2012; Hwang et al., 2014; Jang et al., 2017; Lee et al., 2022) etc in many *in silico, in vitro* and *in vivo* studies.

**Figure 5:**
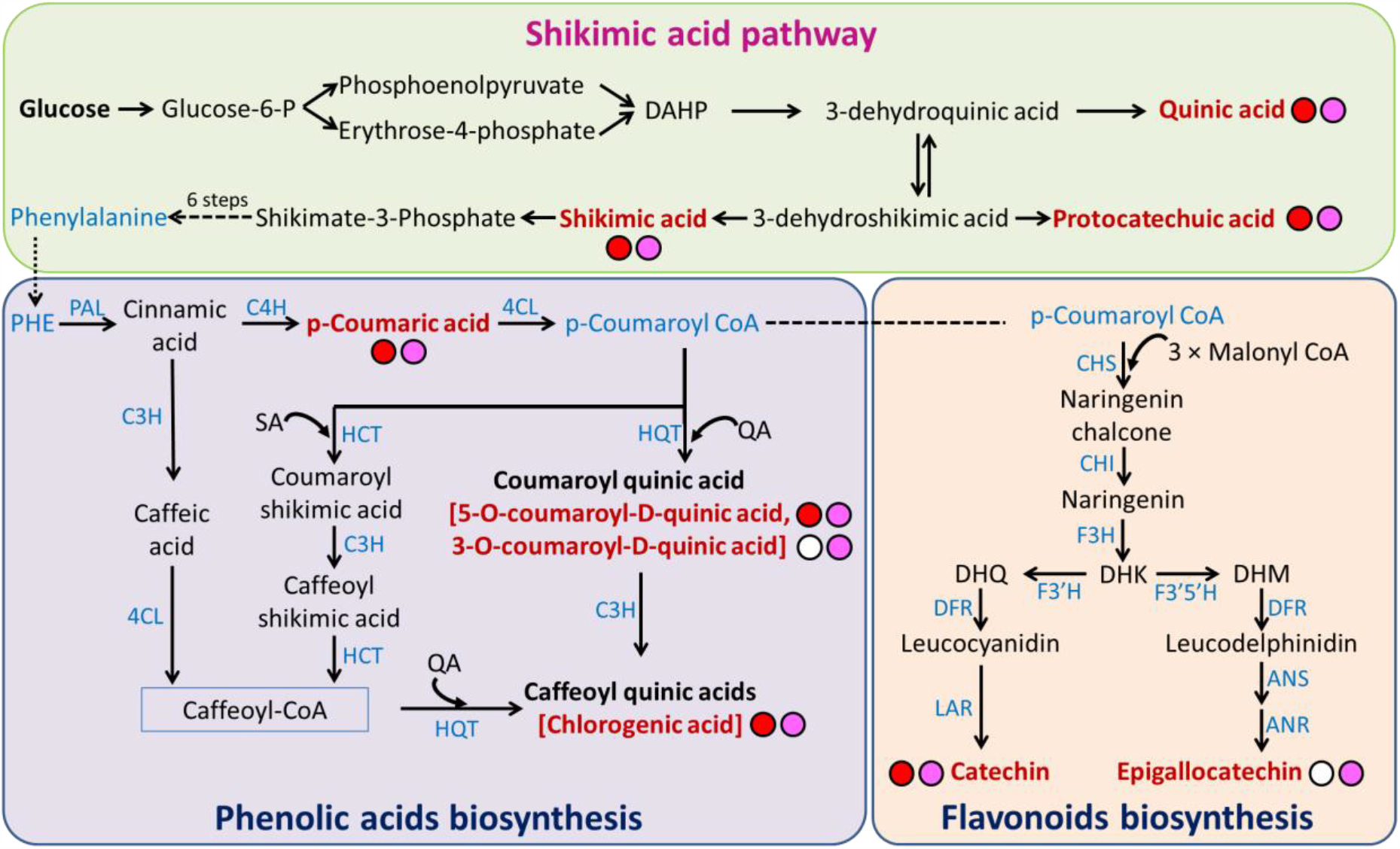
Representation of the secondary metabolite pathways present in *Rhododendron arboreum* and *Rhododendron campanulatum* species highlighted the presence of shikimic acid, phenolic acid biosynthesis and flavonoids biosynthesis pathways. [The phytochemicals identified in the Rhododendron species are in bold letters with maroon color. Red circle – phytochemicals presence in *R. arboreum*; Pink circle -phytochemicals presence in *R. campanulatum* and white circle – absence of phytochemicals. Abbreviations used – DAHP: 3-deoxy-D-arabino-heptulosonate-7-phosphate, PHE: Phenylalanine, PAL: Phenylalanine ammonia-lyase, C4H: Cinnamate 4-hydroxylase, 4CL: 4-coumarate-CoA ligase, C3H: p-coumarate 3′-hydroxilase, HCT: Hydroxycinnamoyl-CoA shikimate/quinate hydroxycinnamoyltransferase, HQT: Hydroxycinnamoyl-CoA quinate hydroxycinnamoyltransferase, QA: Quinic acid, SA: Shikimic acid, CHS: Chalcone synthase, CHI: Chalcone isomerase, F3H: Flavanone 3-hydroxylase, DHK: Dihydrokaempferol, F3’H: Flavonoid 3’-hydroxylase, F3’5’H: Flavonoid 3’,5’-hydroxylase, DHM: Dihydromyricetin, DHQ: Dihydroquercetin, DFR: Dihydroflavonol reductase, LAR: Leucoanthocyanidin reductase, ANS: Anthocyanidin synthase, ANR: Anthocyanidin reductase]

## 4. Conclusion

In this study, the comprehensive profiling of hot aqueous petals extract of *Rhododendron arboreum* and *Rhododendron campanulatum* using multi-analytical platforms was studied. The hot aqueous extract of both species was subjected to LC-ESI-QTOF-MS/MS, GC-MS, and ^1^H-NMR to look into their qualitative and quantitative phytochemical analysis. The phytochemical analysis highlighted the presence of various bioactive metabolites belonging to the phenolic acids and flavonoid classes. Quinic acid, chlorogenic acid, protocatechuic acid, shikimic acid, p-coumaric acid, protocatechuic acid, coumaroyl quinic acids, catechin and epigallocatechin are the key identified metabolites. These bioactive metabolites are already reported to have potential anti-viral, anti-cancerous, anti-diabetic, and anti-inflammatory properties. Profiling of petals extract showed the difference between these two species qualitatively and quantitatively. This was the first study that showed the comprehensive profiling of pink Rhododendron species (*R. campanulatum*) and highlighted the presence of Quinic acid and its derivatives in the flower’s petals. This work is a ready reference for the phytochemical profiles of the Rhododendron petals.

## Acknowledgment

The authors acknowledge the School of Biosciences and Bioengineering (SBB) at IIT Mandi for the GC-MS facilities. Advanced Material Research Centre (AMRC) at IIT Mandi is acknowledged for the LC-ESI-QTOF-MS/MS and NMR facilities. Shagun S thanks CSIR-UGC (09/1058(0024)/2020-EMR-I) for the Ph.D. fellowship.

