## Supplementary material for "Mass spectrometry and NMR spectroscopy profiles of red and pink Rhododendron flower petals establish them as rich sources of bioactive secondary metabolites"

**Supplementary data**

### Comprehensive phytochemical profiling of *R. arboreum* and *R. campanulatum* petals using Mass Spectrometry and Nuclear Magnetic Resonance Spectroscopy

Shagun Shagun[a], Maneesh Lingwan[a], Shyam Kumar Masakapalli[a]*

[a]School of Biosciences and Bioengineering, Indian Institute of Technology Mandi, Kamand 175075, India

| Supplementary | Explanation |
|---|---|
| **Figures** | |
| Supplementary Figure 1 | LC-MS/MS chromatograms of (A) *R. arboreum* (B) *R. campanulatum* petals hot aqueous extract detected in negative ion mode. |
| Supplementary Figure 2 | LC-MS/MS chromatograms of (A) *R. arboreum* (B) *R. campanulatum* petals hot aqueous extract and (C) Quinic acid standard detected in negative ion mode. |
| Supplementary Figure 3 | LC-MS/MS chromatograms of (A) *R. arboreum* (B) *R. campanulatum* petals hot aqueous extract and (C) Chlorogenic acid standard detected in negative ion mode. |
| Supplementary Figure 4 | $^{1}$H-NMR spectra of (A) *R. arboreum* (B) *R. campanulatum* petals hot aqueous extract and (C) Quinic acid standard. |
| Supplementary Figure 5 | $^{1}$H-NMR spectra of (A) *R. arboreum* (B) *R. campanulatum* petals hot aqueous extract and (C) Chlorogenic acid standard. |
| Supplementary Figure 6 | The difference in the abundances of (A) Organic acid and (B) unknowns characterized through LC-MS/MS in *R. arboreum* and *R. campanulatum* petals hot aqueous extracts. |
| Supplementary Figure 7 | Multivariate statistical analysis showed the distinct phytochemicals profiles of *R. arboreum* and *R. campanulatum*. |
| **Tables** | |
| Supplementary Table 1 | LC-ESI-QTOF-MS/MS characteristics of the phytochemicals identified from *Rhododendron arboreum* and *Rhododendron campanulatum* petals hot aqueous extract. |
| Supplementary Table 2 | Phytochemical profiling of *Rhododendron arboreum* petals hot aqueous extracts analysed using GC-MS. |
| Supplementary Table 3 | Phytochemical profiling of *Rhododendron campanulatum* petals hot aqueous extracts analysed using GC-MS. |

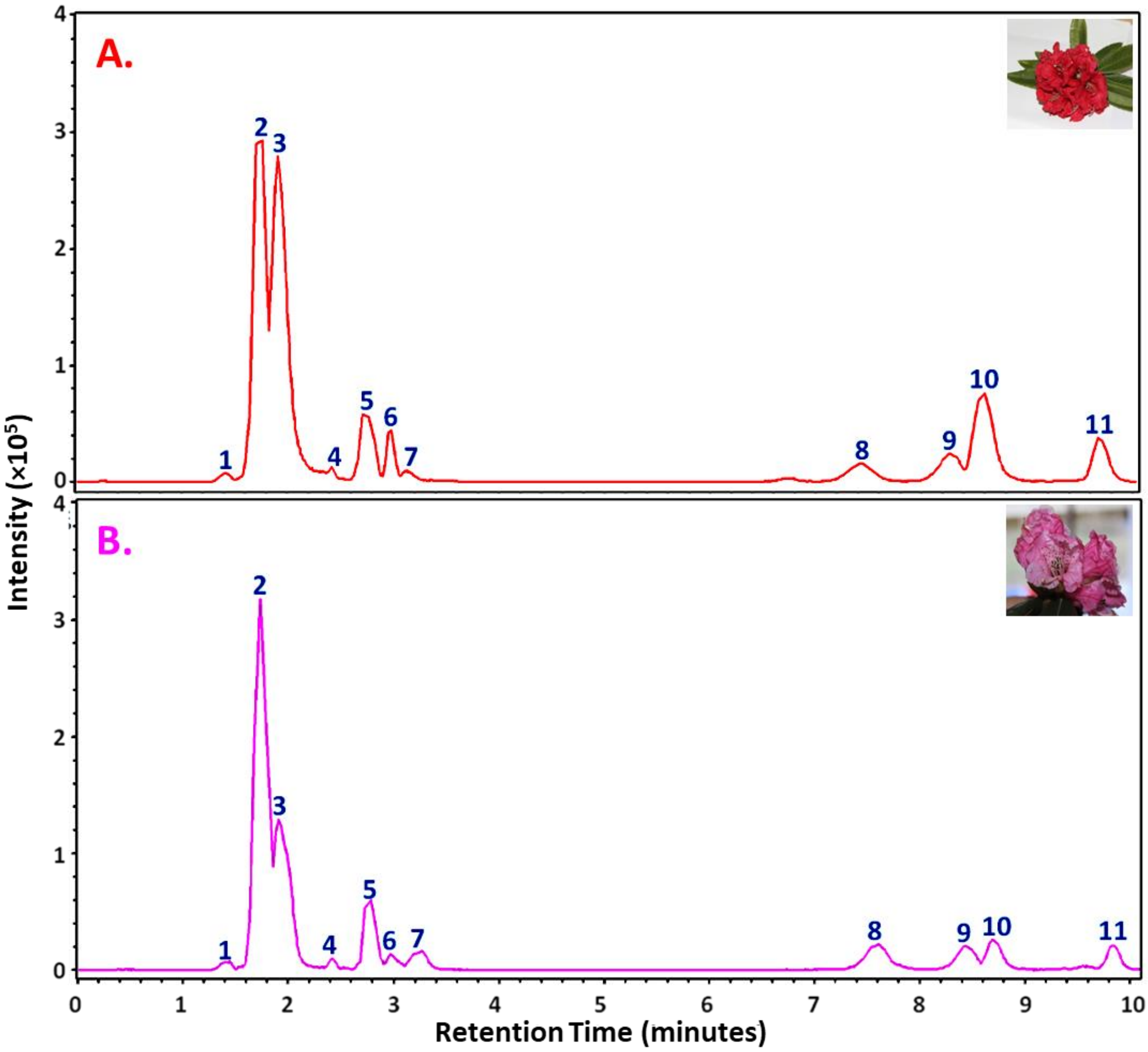


**Supplementary Figure 1:** LC-MS/MS chromatograms of **A.** *Rhododendron arboreum* **B.** *Rhododendron campanulatum* petals hot aqueous extract detected in negative ion mode. Phytochemicals identified are quinic acid (2), malic acid (3), chlorogenic acid (5), protocatechuic acid (6), catechin (8), and coumaroyl quinic acid (10 &11). Peaks 1, 4, 7, and 9 were labeled as unknowns.

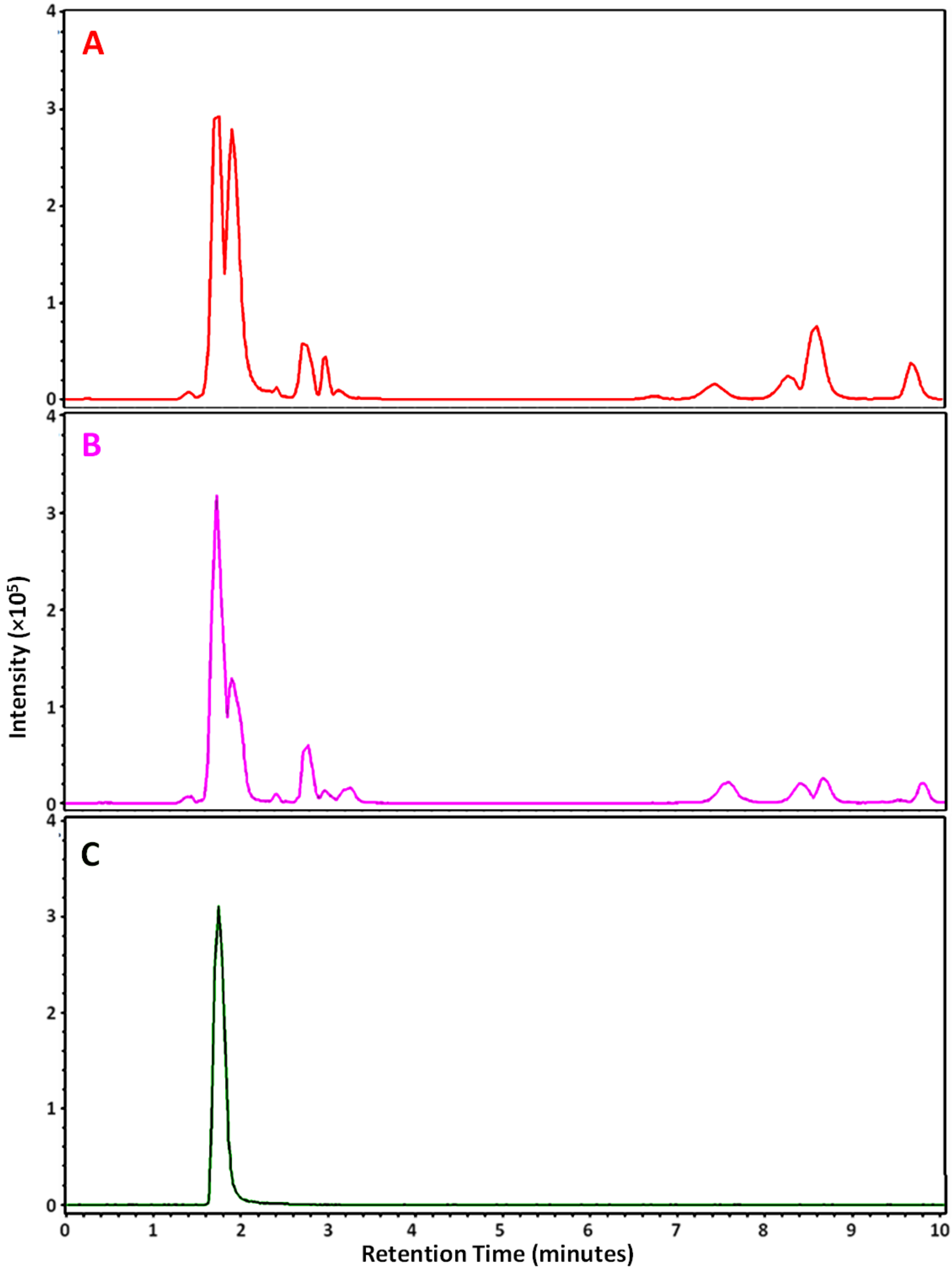


**Supplementary figure 2:** LC-MS/MS chromatograms of **(A)** *Rhododendron arboreum* **(B)** *Rhododendron campanulatum* petals hot aqueous extract and **(C)** Quinic acid standard detected in negative ion mode which confirmed the presence of Quinic acid (retention time-1.8 minutes) in Rhododendron petals.

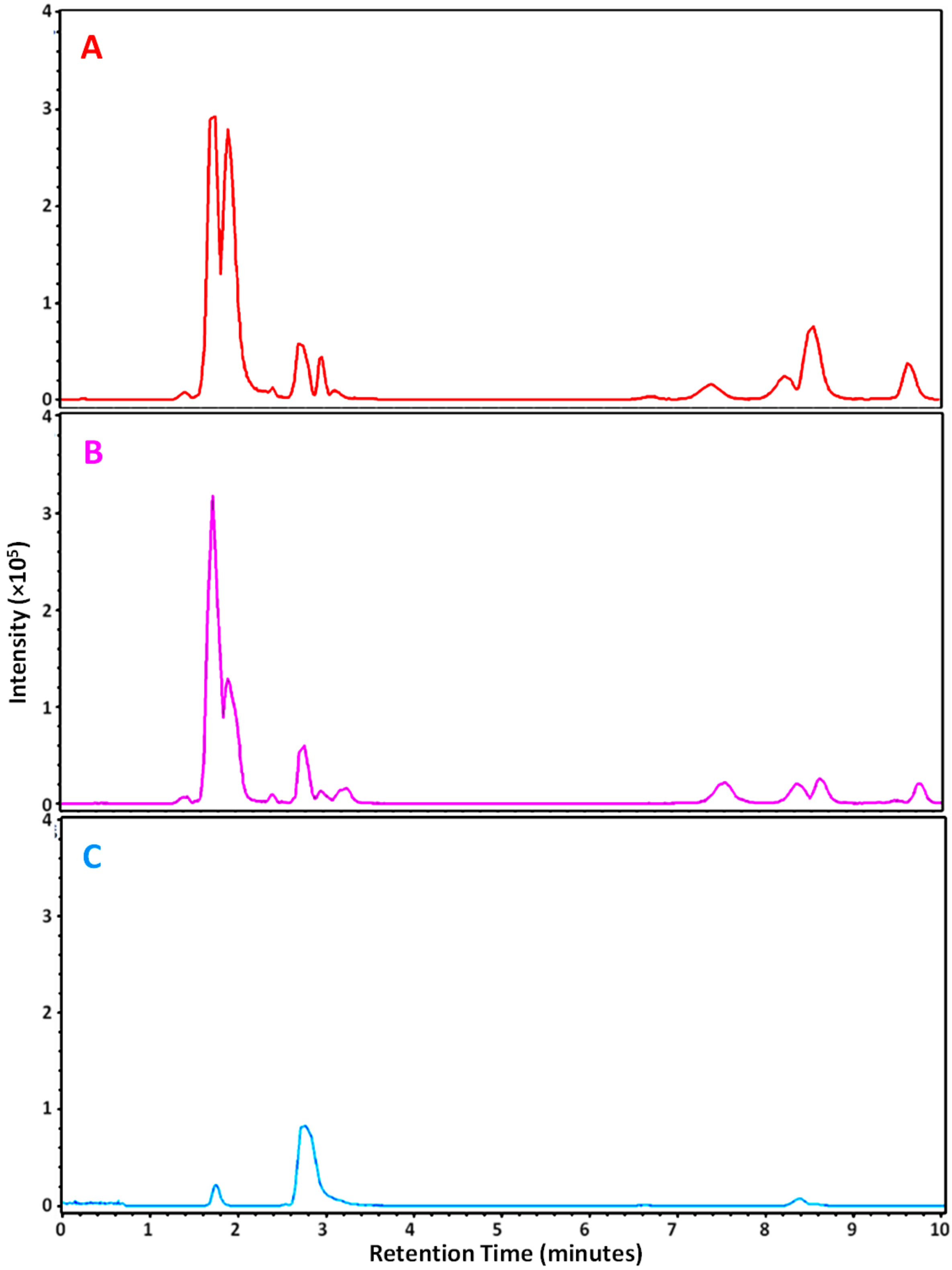


**Supplementary figure 3:** LC-MS/MS chromatograms of **(A)** *Rhododendron arboreum* **(B)** *Rhododendron campanulatum* petals hot aqueous extract and **(C)** Chlorogenic acid standard detected in negative ion mode which confirmed the presence of Chlorogenic acid (retention time-2.8 minutes) in Rhododendron petals.

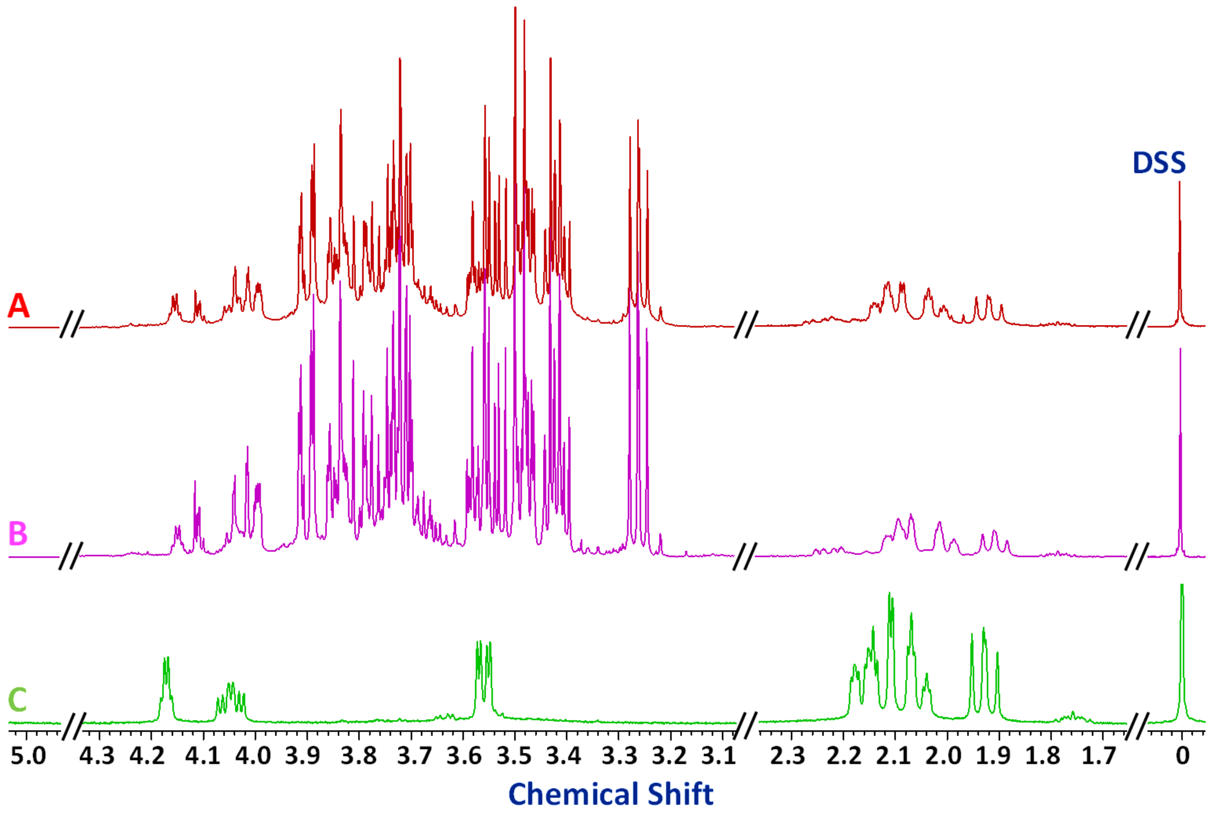


**Supplementary figure 4:** $^{1}$H-NMR spectra of **(A)** *Rhododendron arboreum* **(B)** *Rhododendron campanulatum* petals hot aqueous extract and **(C)** Quinic acid standard which confirmed the presence of Quinic acid in Rhododendron petals.

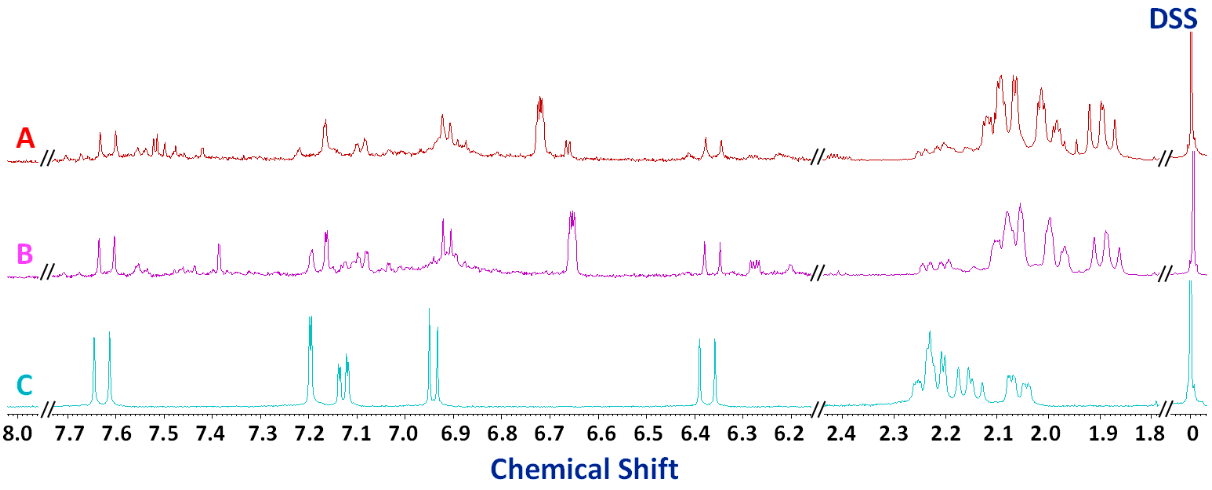


**Supplementary figure 5:** $^{1}$H-NMR spectra of **(A)** *Rhododendron arboreum* **(B)** *Rhododendron campanulatum* petals hot aqueous extract and **(C)** Chlorogenic acid standard which confirmed the presence of Chlorogenic acid in the Rhododendron petals.

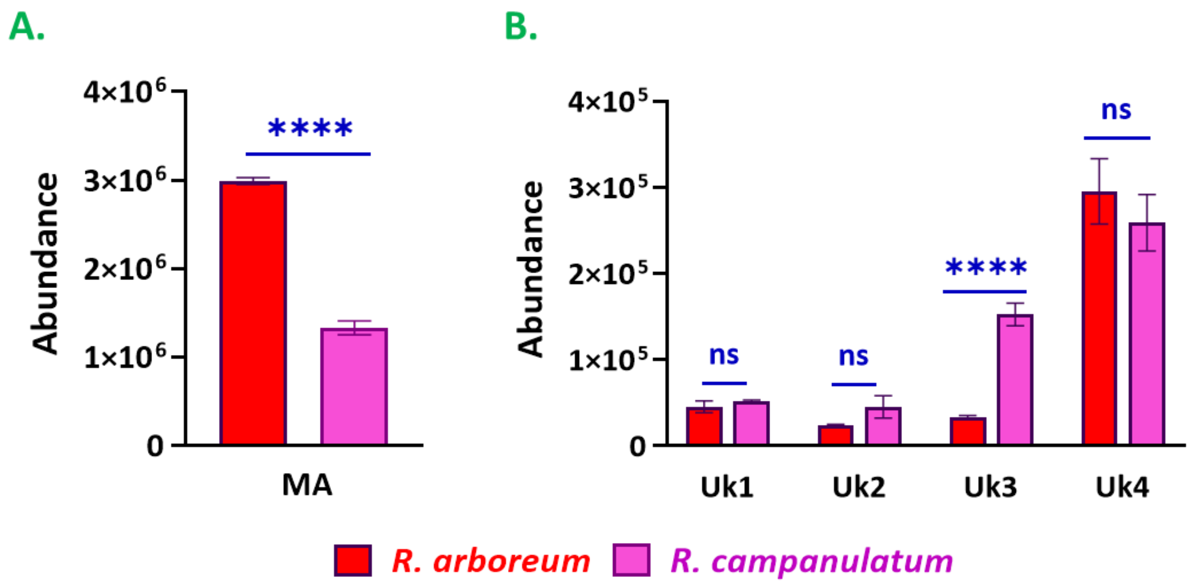


**Supplementary figure 6:** The difference in the abundances of (A) Organic acid, and (B) Unknowns [expressed as Mean ± Standard deviation (n=3)] characterized in *R. arboreum* and *R. campanulatum* petals hot aqueous extracts showed the significant differences (at a significance level of $p < 0.05$) in their levels among both the species. (Abbreviations used: MA-Malic acid, Uk1-Unknown 1; Uk2-Unknown 2; Uk3-Unknown 3; Uk4-Unknown 4; ns-not significant)

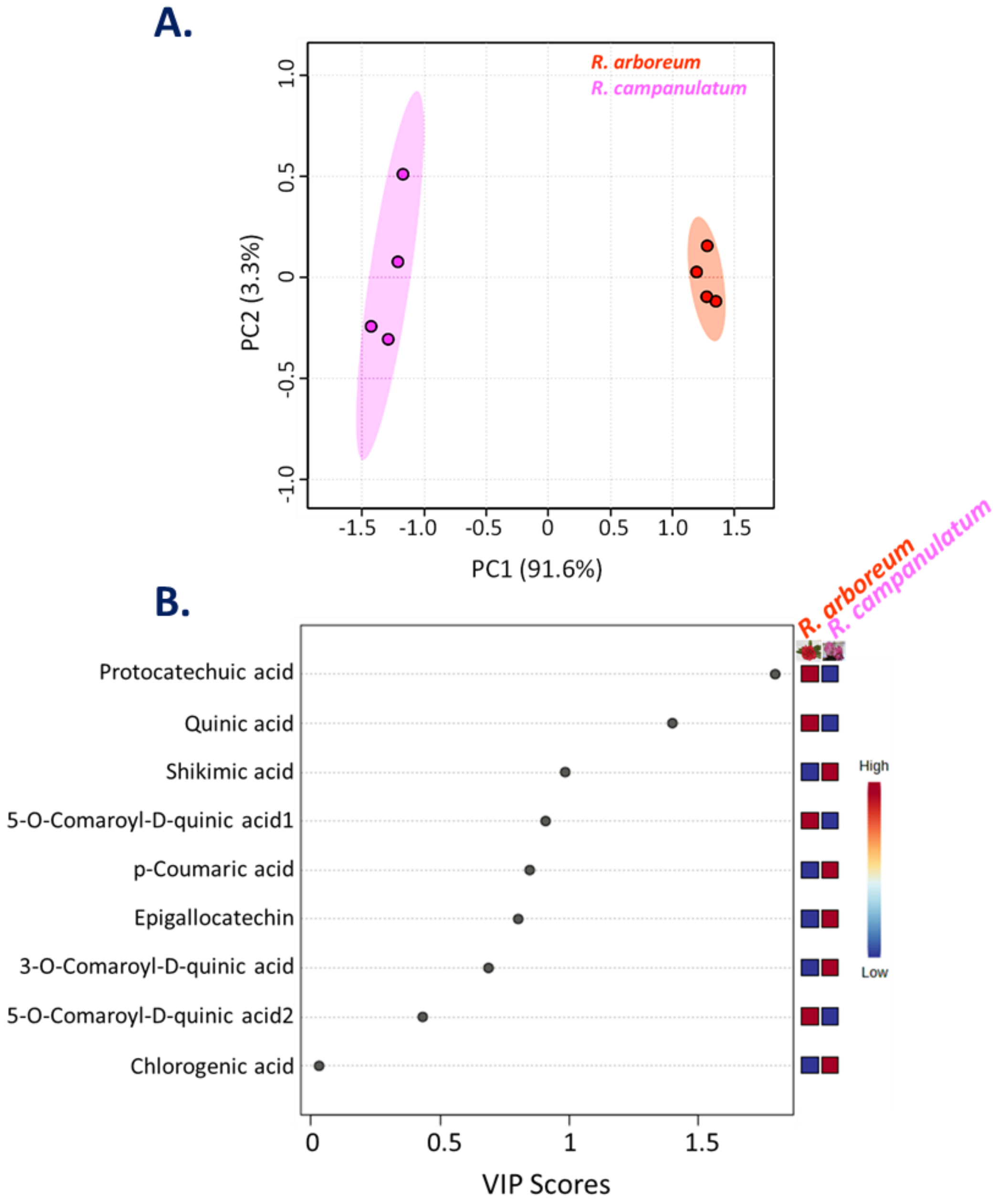


**Supplementary figure 7:** Multivariate statistical analysis showed the distinct phytochemicals profiles of *R. arboreum* and *R. campanulatum*. **A.** PCA (Principal component Analysis) plot indicated the variations among both species. **B.** The variability of the secondary metabolites is presented. Blue and red colored scale is used to showcase the low to the high contribution of phytochemicals.

**Supplementary Table 1.** LC-ESI-QTOF-MS/MS characteristics of the phytochemicals identified from *Rhododendron arboreum* and *Rhododendron campanulatum* petals hot aqueous extract.

| S.No. | Metabolites Identified | Retention time | Chemical formula | $[M-H]^-$ m/z | Fragment ions m/z (MS/MS) | Metabolites class |
|---|---|---|---|---|---|---|
| **1** | Unknown metabolite 1 | 1.4 | Nk* | 273 | $MS^2$ [273]: 115, 131, 159 | Nk* |
| **2** | Quinic acid | 1.8 | $C_7H_{12}O_6$ | 191 | $MS^2$ [191]: 108, 109, 127 | Phenolic acid |
| **3** | Malic acid | 1.9 | $C_6H_8O_7$ | 133 | $MS^2$ [133]: 115 | Organic acid |
| **4** | Unknown metabolite 2 | 2.4 | Nk* | 161 | $MS^2$ [161]: 106 | Nk* |
| **5** | Chlorogenic acid | 2.8 | $C_{16}H_{18}O_9$ | 353 | $MS^2$ [353]: 191 | Phenolic acid |
| **6** | Protocatechuic acid | 3.0 | $C_7H_6O_4$ | 153 | $MS^2$ [153]: 108, 109 | Phenolic acid |
| **7** | Unknown metabolite 3 | 3.1 | Nk* | 353 | $MS^2$ [353]: 191 | Nk* |
| **8** | Catechin | 7.4 | $C_{15}H_{14}O_6$ | 289 | $MS^2$ [289]: 123, 109, 151 | Flavonoid |
| **9** | Unknown metabolite 4 | 8.3 | Nk* | 863 | $MS^2$ [863]: 411, 285, 289 | Nk* |
| **10** | Coumaroyl quinic acid | 8.6 | $C_{16}H_{18}O_8$ | 337 | $MS^2$ [337]: 191, 119 | Phenolic acid |
| **11** | Coumaroyl quinic acid | 9.7 | $C_{16}H_{18}O_8$ | 337 | $MS^2$ [337]: 191, 119 | Phenolic acid |
| Nk*-Not known | | | | | | |

**Supplementary Table 2.** Phytochemical profiling of *Rhododendron arboreum* petals hot aqueous extracts analysed using GC-MS. The retention time (RT), m/z product ion, relative peak area [mean ± standard deviation (n=4)] and probability scores against the NIST17 are presented. Ribitol (0.01%) was used as internal standard.

| S. No. | Phytochemicals | RT | % Area | Probability score* (m/z) |
|---|---|---|---|---|
| 1 | 2,3-Butanediol | 11.40 | 0.3 ± 0.0 | 71 (147, 133, 117) |
| 2 | 2,3-Butanediol | 11.54 | 0.7 ± 0.0 | 74 (147, 133, 117) |
| 3 | Lactic Acid | 12.05 | 0.5 ± 0.1 | 84 (147, 191, 117) |
| 4 | Glycolic acid | 12.32 | 0.4 ± 0.0 | 86 (177, 205, 147) |
| 5 | Ethanolamine | 15.08 | 0.1 ± 0.0 | 70 (174, 147, 133) |
| 6 | Maleic acid | 15.64 | 1.0 ± 0.1 | 88 (245, 147, 133) |
| 7 | Succinic acid | 15.79 | 1.4 ± 0.1 | 76 (247, 147, 129) |
| 8 | Glyceric acid | 15.95 | 1.1 ± 0.0 | 86 (189, 292, 147) |
| 9 | Fumaric acid | 16.28 | 0.3 ± 0.0 | 95 (245, 147, 143) |
| 10 | D- (-)-Citramalic acid | 17.85 | 0.1 ± 0.0 | 91 (247, 349, 203) |
| 11 | Malic acid | 18.09 | 17.5 ± 0.3 | 98 (233, 245, 189) |
| 12 | 2,3,4 Tri hydroxybutyric acid | 18.89 | 1.2 ± 0.0 | 91 (292, 220, 205) |
| 13 | 3-Hydroxy-3-methylglutaric acid | 19.45 | 0.8 ± 0.3 | 97 (247, 363, 273) |
| 14 | Tartaric acid | 19.85 | 1.5 ± 0.1 | 88 (292, 219, 147) |
| 15 | Shikimic acid | 22.11 | 3.5 ± 0.3 | 86 (204, 255, 147) |
| 16 | Citric acid | 22.34 | 6.1 ± 0.8 | 91 (273, 363, 465) |
| 17 | Protocatechuic acid | 22.40 | 3.6 ± 0.1 | 90 (193, 370, 311) |
| 18 | Quininic acid | 23.03 | 40.3 ± 1.7 | 93 (345, 255, 191) |
| 19 | p-Coumaric acid | 25.08 | 0.5 ± 0.0 | 83 (219, 293, 249) |
| 20 | D-Gluconic acid | 26.27 | 4.8 ± 0.2 | 80 (333, 292, 205) |
| 21 | Galactaric acid | 27.58 | 0.5 ± 0.1 | 76 (333, 292, 189) |
| 22 | Palmitic Acid | 28.00 | 2.1 ± 0.1 | 98 (133, 145, 129) |
| 23 | Stearic acid | 34.84 | 0.9 ± 0.1 | 92 (341, 145, 132) |
| 24 | 1-Monopalmitin | 42.42 | 3.0 ± 0.2 | 97 (371, 459, 313) |
| 25 | 5-O-Coumaroyl-D-quinic acid | 46.65 | 1.6 ± 0.1 | 88 (345, 219, 255) |
| 26 | 5-O-Coumaroyl-D-quinic acid | 48.48 | 3.6 ± 0.5 | 93 (345, 219, 255) |
| 27 | Chlorogenic acid | 50.31 | 2.9 ± 0.4 | 95 (345, 255, 307) |
| *- Match score with NIST library | | | | |

**Supplementary Table 3.** Phytochemical profiling of *Rhododendron campanulatum* petals hot aqueous extracts analysed using GC-MS. The retention time (RT), m/z product ion, relative peak area [mean ± standard deviation (n=4)] and probability scores against the NIST17 are presented. Ribitol (0.01%) was used as internal standard.

| S. No. | Phytochemicals | RT | % Area | | | Probability score* (m/z) | |
|---|---|---|---|---|---|---|---|
| 1 | Boric acid | 10.44 | 0.4 | ± | 0.1 | 79 | (221, 263, 205) |
| 2 | Ethylene glycol | 10.58 | 0.5 | ± | 0.1 | 78 | (191, 147, 133) |
| 3 | 2,3-Butanediol | 11.30 | 0.9 | ± | 0.2 | 73 | (147, 133, 117) |
| 4 | 2,3-Butanediol | 11.43 | 0.7 | ± | 0.1 | 71 | (147, 133, 117) |
| 5 | Lactic Acid | 11.95 | 0.4 | ± | 0.0 | 82 | (147, 191, 117) |
| 6 | n-Butylamine | 12.31 | 0.4 | ± | 0.1 | 85 | (174, 202, 128) |
| 7 | Hydroxylamine | 12.64 | 0.3 | ± | 0.1 | 83 | (249, 204, 147) |
| 8 | Ethanolamine | 14.97 | 1.4 | ± | 0.1 | 79 | (174, 147, 133) |
| 9 | Maleic acid | 15.55 | 0.9 | ± | 0.2 | 70 | (245, 147, 133) |
| 10 | Succinic acid | 15.70 | 0.9 | ± | 0.0 | 70 | (247, 147, 129) |
| 11 | Glyceric acid | 15.85 | 1.5 | ± | 0.1 | 85 | (189, 292, 147) |
| 12 | Fumaric acid | 16.20 | 0.3 | ± | 0.1 | 91 | (245, 147, 143) |
| 13 | D- (-)-Citramalic acid | 17.74 | 0.2 | ± | 0.0 | 80 | (247, 349, 203) |
| 14 | Malic acid | 17.98 | 8.0 | ± | 0.1 | 97 | (233, 245, 189) |
| 15 | 4-Aminobutanoic acid | 18.50 | 2.1 | ± | 0.1 | 95 | (304, 174, 147) |
| 16 | 2,3,4-Tri hydroxybutyric acid | 18.78 | 2.7 | ± | 0.4 | 95 | (292, 220, 205) |
| 17 | 3-Hydroxy-3-methylglutaric acid | 19.35 | 2.1 | ± | 0.2 | 97 | (247, 363, 273) |
| 18 | Tartaric acid | 19.71 | 2.3 | ± | 0.2 | 84 | (292, 219, 147) |
| 19 | Shikimic acid | 21.99 | 13.3 | ± | 0.6 | 88 | (204, 255, 147) |
| 20 | Citric acid | 22.25 | 7.2 | ± | 0.8 | 96 | (273, 363, 465) |
| 21 | Quinic acid | 22.86 | 10.7 | ± | 1.7 | 90 | (345, 255, 191) |
| 22 | p-Coumaric acid | 24.86 | 1.6 | ± | 0.3 | 82 | (219, 293, 249) |
| 23 | D-Gluconic acid | 25.99 | 6.6 | ± | 0.1 | 94 | (333, 292, 205) |
| 24 | D-Gluconic acid | 26.38 | 6.0 | ± | 0.4 | 91 | (333, 292, 205) |
| 25 | Galactaric acid | 27.27 | 4.7 | ± | 0.2 | 73 | (333, 292, 189) |
| 26 | Palmitic Acid | 27.70 | 5.2 | ± | 0.2 | 95 | (133, 145, 129) |
| 27 | Stearic acid | 34.61 | 2.9 | ± | 0.2 | 92 | (341, 145, 132) |
| 28 | 1-Monomyristin | 38.41 | 0.1 | ± | 0.0 | 88 | (343, 431, 285) |
| 29 | 1-Monopalmitin | 42.22 | 3.1 | ± | 0.1 | 96 | (371, 459, 313) |
| 30 | Glycerol monostearate | 45.43 | 3.4 | ± | 0.1 | 97 | (399, 487, 267) |
| 31 | 5-O-Coumaroyl-D-quinic acid | 46.45 | 2.0 | ± | 0.1 | 90 | (345, 219, 255) |
| 32 | Epigallocatechin | 47.46 | 0.0 | ± | 0.0 | 86 | (456, 355, 280) |
| 33 | 5-O-Coumaroyl-D-quinic acid | 48.28 | 2.6 | ± | 0.1 | 94 | (345, 219, 255) |
| 34 | 3-O-Coumaroyl-D-quinic acid | 49.51 | 0.0 | ± | 0.0 | 78 | (345, 219, 255) |
| 35 | Chlorogenic acid | 50.09 | 4.5 | ± | 0.2 | 96 | (345, 255, 307) |
| 36 | Trisdibutylphenyl phosphite | 54.74 | 0.1 | ± | 0.0 | 88 | (441, 308, 147) |
| *- Match score with NIST library | | | | | | | |
